# Origin and evolutionary landscape of *Nr2f* transcription factors across Metazoa

**DOI:** 10.1101/2021.07.22.453436

**Authors:** Ugo Coppola, Joshua S. Waxman

**Affiliations:** Molecular Cardiovascular Biology Division and Heart Institute, Cincinnati Children’s Research Foundation, Cincinnati, OH 45229, USA; Department of Pediatrics, University of Cincinnati College of Medicine, Cincinnati, OH 45229, USA

## Abstract

**Background:** Nuclear Receptor Subfamily 2 Group F (Nr2f) orphan nuclear hormone transcription factors (TFs) are fundamental regulators of many developmental processes in invertebrates and vertebrates. Despite the importance of these TFs throughout metazoan development, previous work has not clearly outlined their evolutionary history.

**Results:** We integrated molecular phylogeny with comparisons of intron/exon structure, domain architecture, and syntenic conservation to define critical evolutionary events that distinguish the *Nr2f* gene family in Metazoa. Our data indicate that a single ancestral pre-metazoan *Nr2f* gene, we have termed *Nr2f1/2/5/6,* predated six main Bilateria subfamilies, which include a single *Nr2f1/2/5* homolog that is present throughout protostomes and invertebrate deuterostomes, *Nr2f1/2* homologs in agnathans, and *Nr2f1*, *Nr2f2*, *Nr2f5*, *Nr2f6* orthologs that are found in gnathostomes. The three *Nr2f1/2* members in agnathans are due to independent expansions not found in gnathostomes, while the vertebrate *Nr2f1*, *Nr2f2*, *Nr2f5* members arose from whole-genome duplications (WGDs). However, *Nr2f6* members are the most divergent subfamily, likely originating from an ancient duplication, and are only retained by gnathostomes. Interestingly, *Nr2f5* TFs have been independently lost in both cartilaginous fish and amniotes, such as humans. Furthermore, our analysis shows there are differential expansions and losses of *Nr2f* genes in teleosts following their additional rounds of WGDs.

**Conclusion:** Overall, our evolutionary genomic analysis of Nr2f proteins helps to reveal the origins and previously unrecognized relationships of this ancient transcription factor family, which may allow for greater insights into the conservation of Nr2f functions that shape Metazoan body plans.

## Introduction

Nuclear hormone receptors (NRs) form a large, ancient superfamily of transcription factors (TFs) found in all Metazoa [1]. While NR functions are often dictated by interactions with specific ligands, including steroids, thyroid hormones, and retinoids [2, 3], the ligands for many NRs, called orphan NRs, are still not known [4]. Nuclear Receptor Subfamily 2 Group F Members (Nr2fs), initially named Chicken ovalbumin upstream promoter-transcription factors (Coup-TFs) due to their ability to bind the COUP element of the ovalbumin gene [5–7], are some of the most highly studied orphan NRs. Despite an overall expansion of the NR superfamily [1, 2], invertebrate phyla appear to have predominantly retained a single *Nr2f* gene. Only one *Nr2f* member is present in the protostome *Drosophila melanogaster* (fly), early-branching deuterostome *Strongylocentrotus purpuratus* (sea urchin) [8, 9], and invertebrate chordates *Branchiostoma floridae* (amphioxus) and *Ciona robusta* (sea squirt) [10, 11]. However, the number of *Nr2f* genes in early-branching metazoans is presently less clear. In cnidarians, one *Nr2f* has been reported in *Hydra*, while multiple have been reported in *Nematostella* [12, 13]. In contrast to most invertebrates, vertebrates have exhibited a significant expansion of the *Nr2f* family, with gnathostomes having multiple *Nr2f* genes. Furthermore, teleosts possess additional *Nr2f* Ohnologs (duplicates originating from whole-genome duplication (WGD)) [14], most likely reflecting the additional WGDs that have occurred in the teleost lineage [15, 16].

Nr2f proteins are highly conserved at the sequence level throughout Metazoa (Giguere, 1999). From the N-terminus to the C-terminus, all Nr2f proteins have six domains (Fig 1): an A/B domain, which contains the activating function-1 (AF-1) domain; the C domain, which contains the DNA-binding domain (DBD); the D domain (a linker); the E domain, which is comprised of the ligand-binding domain (LBD) and an AF-2 domain; the F domain (C-terminal) [17]. While the A/B domains are the most divergent in sequence, strikingly, the DBDs and LBDs of Nr2f members even from distantly related species (e.g. fly, sea urchin, frog, zebrafish, mouse, and human) are ∼94% identical [18]. The extremely high degree of conservation among several species implies the preservation of critical roles for Nr2f in development and differentiation [18, 19]. Requirements for *Nr2f* genes have been found in organs of all three germ layers during embryogenesis [12, 19]. For instance, the *Drosophila Nr2f* homolog, called *seven up* (*svp*) is required for retinal, dorsal vessel, and liver development [20, 21]. Furthermore, Nr2f transcription factors in vertebrates appear to both have acquired diverse and retained redundant functions. For instance, in mice, *Nr2f1* is predominantly required for neural development with a role in regulation of premigratory and migratory neural crest cells in the developing hindbrain [22, 23]. However, the mouse *Nr2f2* gene is required for differentiation of mesodermal derivatives, including atrial cardiomyocytes of the heart and venous endothelial cells [19,24,25]. An example of redundancy are zebrafish *nr2f1a* and *nr2f2*, which are both required for proper ventricular cardiomyocyte and cranial muscle specification [26].

**Fig 1.**
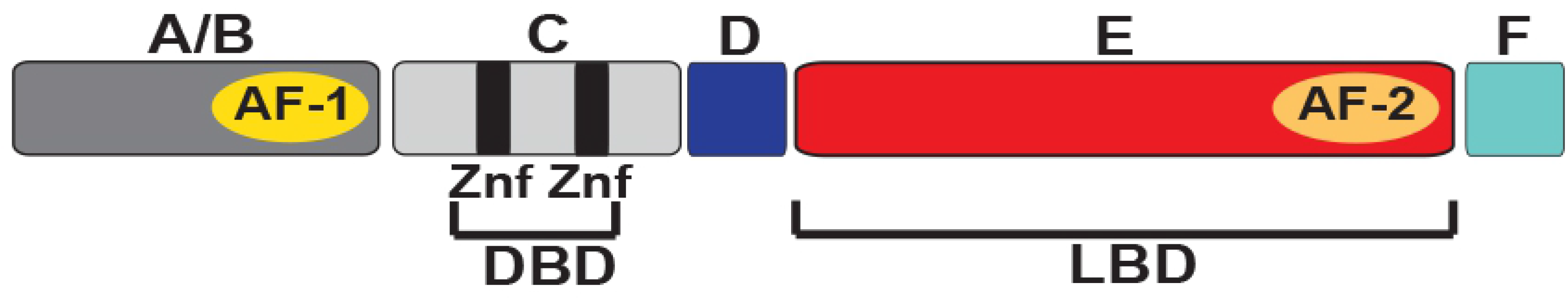
Schematic of conserved domain architecture of Nr2f transcription factors. A/B (N-terminal variable domain with transactivating AF-1 domain), C (DBD, which two contains two Zinc finger (Znf) motifs), D (a linker domain), E (LBD plus transactivating AF-2 domain), and F (C-terminal).

While Nr2f proteins were initially identified as transcriptional activators of chicken ovalbumin gene [5], they have since been shown to function directly as both transcriptional activators and repressors in several developmental contexts [19,27,28]. Nr2fs can bind a range of different response elements [29–31] and in signaling reporter assays can compete with and inhibit retinoic acid receptors [32]. *In vivo* they bind numerous targets that reflect their various requirements in the specific tissues. For instance, *Nr2f1* KO mice also have inner ear defects [33]. In the mouse inner ear, direct targets of Nr2f1 include fatty acid binding protein 7 (FABP7), cellular retinoic acid binding protein 1 (CRABP1)[34], microRNA-140 (mi-R140), and Kruppel-like 9 (Klf9) [34, 35]. In adipogenesis, Nr2f2 directly represses peroxisome proliferator-activated receptor γ (PPARγ) downstream of canonical Wnt/β-catenin signaling [36]. In the mammalian heart, Nr2f2 is thought to directly orchestrate a regulatory network that facilitates atrial cardiomyocyte identity through concurrently promoting *Tbx5* and repressing *Irx4* and *Hey2*, the latter of which promote ventricular cardiomyocyte identity [37]. Thus, Nr2fs can activate and repress a range of direct targets related to their functions in specific tissues.

Despite the conservation and clear importance of this gene family to numerous developmental processes in Metazoa, we still do not completely understand the evolution of Nr2f TFs. Here, we investigated *Nr2f* family evolution through a combination of phylogenetic, domain architecture, intron/exon structure, and genomic synteny analyses. Our data show that the single *Nr2f1/2/5/6* gene in cnidarians and placozoans, represents the ancestral *Nr2f* to those found in protostomes and deuterostomes. Importantly, a single *Nr2f* homolog, which we have named *Nr2f1/2/5*, is present in the majority of invertebrates, while most vertebrate genomes contain *Nr2f1*, *Nr2f2, Nr2f5* and *Nr2f6* orthologs. Interestingly, the invertebrate *Nr2f1/2/5* and agnathan *Nr2f1/2* homologs have retained the greatest similarity with vertebrate *Nr2f1* and *Nr2f2* paralogs, while *Nr2f5* has been independently lost in the lineages of cartilaginous fish and amniotes. The vertebrate-specific *Nr2f6* subfamily, the most divergent *Nr2f* subfamily, predates the vertebrate radiation, suggesting a different evolutionary relationship to the other *Nr2f* genes. Overall, our data clarify the relationships among *Nr2f* genes within Metazoa and define the expansion, divergence, and independent loss events of extant *Nr2f* genes in vertebrates, which will allow us to make meaningful inferences about the conserved developmental functions of this family that have helped mold animal body plans.

## Results

### Phylogenetic reconstruction of *Nr2f* evolution in animals

Although previous work has investigated the homology of some Nr2fs within metazoans, these analyses were primarily focused on their relationship to other NRs and were limited by the comparatively little genomic information at the time [1,12,38]. To garner a better understanding of how the *Nr2f* family has evolved in animals, we performed a phylogenetic analysis using 130 Nr2f proteins with representatives from placozoans to mammals (Fig 2). Pre-metazoan models *Amphimedon queenslandica* (sponge) and *Mnemiopsis leidyi* (ctenophore) were not included, as we did not find putative *Nr2f* orthologs based on current databases. A maximum-likelihood (ML) phylogenetic tree showed the existence of distinct *Nr2f* subfamilies (Fig 2), which were also supported using a Bayesian model selection (S1 Fig). The placozoan *Trichoplax adhaerens* and cnidarian *Nematostella vectensis* Nr2fs, which we now call the Nr2f1/2/5/6, were at the root of all the metazoan Nr2f proteins (Fig 2). Interestingly, the tree indicated that the three distinct Nr2f1/2/5/6 members in *N. vectensis* that were previously reported are likely the result of gene duplications [13]. The protostome and deuterostome *Nr2f* sequences clustered into six subfamilies, which we have called *Nr2f1/2/5*, *Nr2f1/2, Nr2f1*, *Nr2f2*, *Nr2f5,* and *Nr2f6*. The *Nr2f1/2/5* subfamily genes, which are highly conserved, yet evolutionary divergent from the *Nr2f1/2/5/6* genes present in early-branching animals, were found in invertebrate protostomes, invertebrate deuterostomes (hemichordates, echinoderms), and invertebrate chordates (amphioxus, tunicates). This Nr2f1/2/5 group is more closely related to the branch that includes Nr2f1/2s from the agnathan vertebrate (lamprey) and vertebrate Nr2f1 and Nr2f2 proteins (Fig 2). The clustering of invertebrate Nr2f1/2/5 proteins with Nr2f1, Nr2f2, and Nr2f5 of gnathostomes suggests these proteins arose from distinct duplicative events during vertebrate evolution. Furthermore, the three agnathan Nr2f proteins (Fig 2), which we have called Nr2f1/2A, Nr2f1/2B, and Nr2f1/2C, diverge and cluster together at the base of the vertebrate-specific Nr2f1 and Nr2f2 proteins (Fig 2), suggesting that the duplications leading to these proteins in agnathans were distinct from those that gave rise to the Nr2f paralogs in gnathostomes.

**Fig 2.**
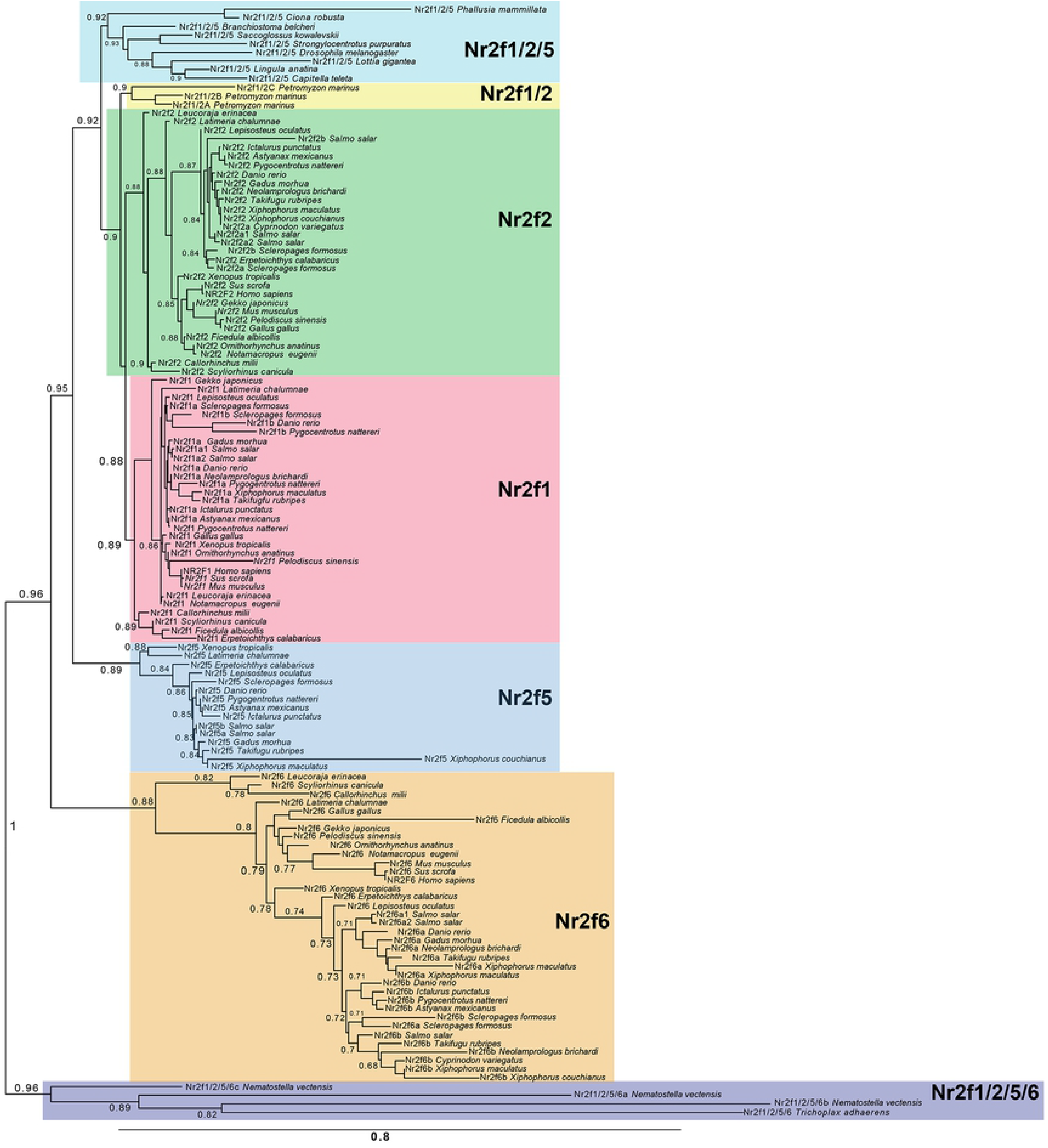
Evolutionary reconstruction of Nr2f proteins in metazoans. Phylogenetic tree of Nr2f members demonstrating the existence of six protein classes: Nr2f1/2/5/6 (violet box), Nr2f1/2/5 (light blue box), Nr2f1/2 (yellow box), Nr2f1 (pink box), Nr2f2 (green box), Nr2f5 (blue box), Nr2f6 (orange box). Values at the branches indicate replicates obtained using the aLRT method.

Our analysis shows that all gnathostomes have retained Nr2f1, Nr2f2 and Nr2f6, whereas Nr2f5 has been lost by cartilaginous fish (chimaerae, sharks, skates) and amniotes (reptiles, avians, mammals), suggesting these were independent losses. Within the vertebrate Nr2f1, Nr2f2, and Nr2f5 cluster, the branching and distances from our phylogenetic tree indicate that Nr2f1 and Nr2f2 are more highly conserved (Fig 2; S1 Fig). Remarkably, Nr2f6 is the most divergent vertebrate Nr2f subfamily and occupies a distinct position in the tree, suggesting a different duplication event for its origin that occurred prior to gnathostomes. Moreover, to analyze the impact of WGDs that took place in teleosts [15, 16], and specifically, in salmonids [39], we surveyed the Nr2f proteins of 12 teleost species (Fig 2). Consistent with the WGDs in these species, there was a tremendous expansion of the *Nr2f* family in this clade, although it was accompanied by differential *Nr2f* paralog losses in some species (Fig 2). To further interrogate the evolution of the Nr2f proteins, we examined alignments of the highly conserved zinc-fingers (Znf) within their DNA-binding domains (DBDs) using representatives from each subfamily (Fig 3). Although there is a high degree of conservation in all the examined Nr2fs, the amino acid changes in the DBDs parallels the phylogenetic results of the whole proteins. The ancient Nr2f1/2/5/6 of early-branching metazoans showed a high degree of variability and multiple differences with respect to Nr2f1/2/5 DBDs of other invertebrates and the Nr2f DBDs in agnathans and gnathostomes. There is high similarity between Nr2f1/2, Nr2f1 and Nr2f2 DBDs in agnathans and gnathostomes, whereas Nr2f5 and Nr2f6 DBDs of gnathostomes exhibited specific changes that are consistent with their positions in the phylogenetic trees (Fig 2; Fig 3). Thus, our phylogenetic reconstruction of *Nr2f* genes in metazoans generally shows the presence of a single ortholog in invertebrates and significant expansion of the family in vertebrates that is punctuated with independent losses of *Nr2f5* in cartilaginous fishes and amniotes.

**Fig 3.**
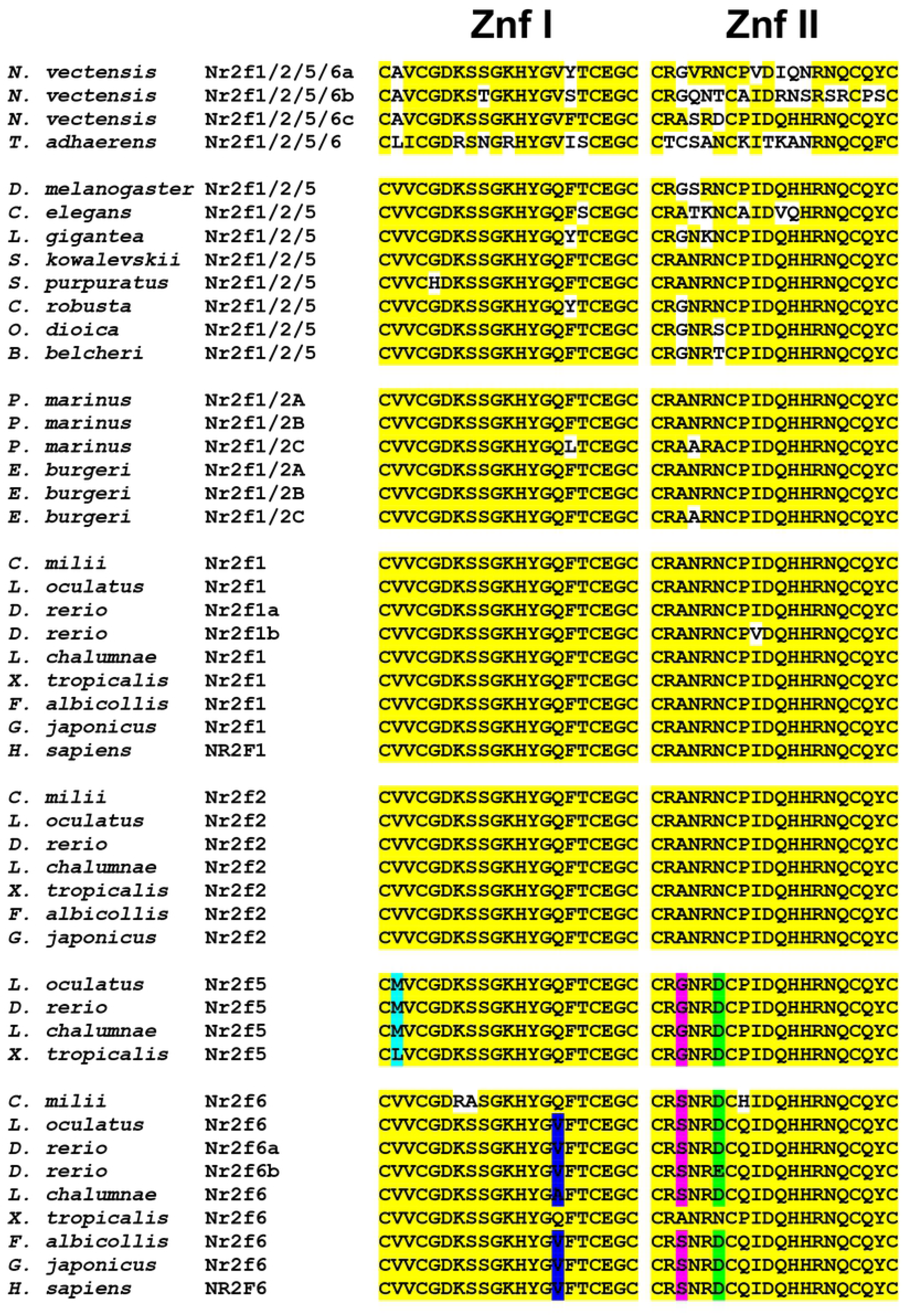
Zinc finger (Znf) motifs within the DBD of the Nr2f family. Alignments of first (I) and second (II) Znfs. Yellow represents conserved amino acids conserved during evolution. White indicates amino acids that are not conserved. Turquoise and blue indicate the amino acid changes that are conserved within Znf I of Nr2f5 and −6, respectively. Magenta and green indicate the amino acid that are conserved within Znf II of Nr2f5 and −6, respectively.

### *Nr2f* genes have conserved intron codes

To complement the phylogenetic analysis of *Nr2f* genes, we first analyzed the conservation of *Nr2f* intron/exon structure [40–42]. Intron/exon junctions from early-branching metazoans and vertebrates matching the transcripts and the translated proteins were mapped and given a score for the intron phases (S1 File), with 0, 1 and 2 introns falling before the first, second and third bases of a codon, respectively. Then, we mapped the introns on a protein alignment comprising the highly conserved Nr2f DBDs and LBDs of Nr2f proteins (S1 File). We found that two “phase 1” introns (one before and one within the LBD) are preserved in all the extant *Nr2f* subfamilies (Fig 4A; S1 File). However, *Nr2f6* genes exhibited a “phase 2” intron inside the second zinc-finger belonging to the nuclear receptor DNA-binding site (Fig 4A; S1 File). The conservation of intron/exon junctions in the examined *Nr2f* genes allows two groups to be distinguished: one constituted by *Nr2f1/2/5/6*, *Nr2f1/2/5*, *Nr2f1/2*, *Nr2f1*, *Nr2f2*, *Nr2f5*, and one comprising only *Nr2f6* (Fig 4B). Thus, our analysis of intron/exon boundaries demonstrates the existence of a highly conserved intron code throughout *Nr2f* family members that also robustly supports the divergence of *Nr2f6* genes.

**Fig 4.**
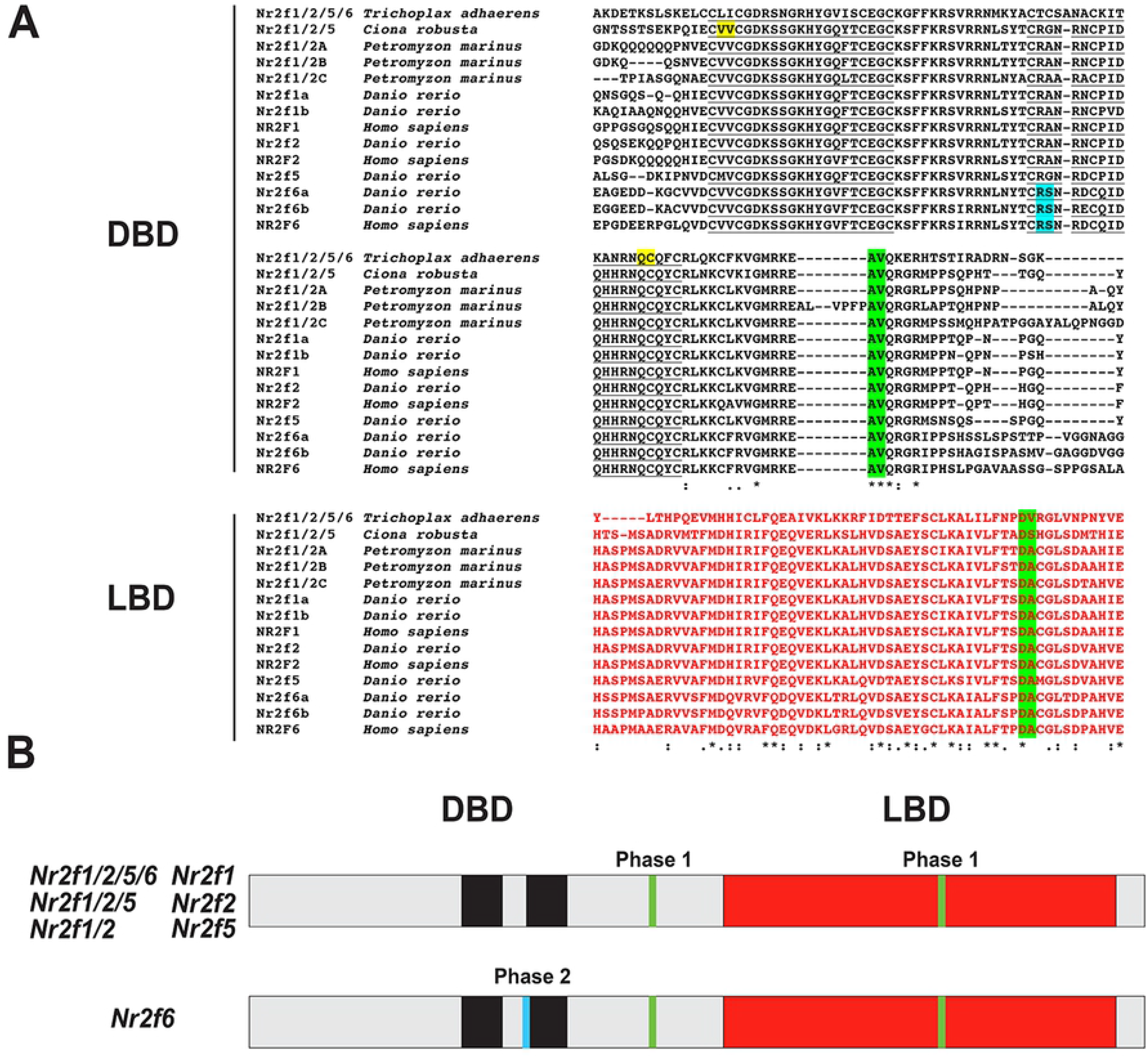
Intron code of the *Nr2f* family in metazoans. (A) Protein alignment showing conservation of intron/exon structures in Nr2f members, with a portion comprising the second DBD (underlined) and LBD (red). Asterisks indicate 100% amino acid conservation. Colons indicate high levels (>90%) amino acid conservation. Periods indicate moderate levels (50-89%) of amino acid conservation. Phase1-introns (green) and the *Nr2f6*-specific Phase 2-intron (turquoise). (B) Schematization of distinct intron/exon structures of *Nr2f1/2/5/6*, *Nr2f1/2/5*, *Nr2f1/2, Nr2f1*, *Nr2f2*, *Nr2f5* genes and of *Nr2f6*. Black boxes represent the zinc-fingers motifs within the DBD, while pink bars indicate the LDB. Small vertical bars were used to portray the ultra-conserved Phase1-introns and the *Nr2f6*-specific Phase 2-intron.

### Synteny analysis defines differential duplications and losses in the *Nr2f* family

In order to confirm the specific homologies indicated from the phylogenetic analysis, we next carried out an extensive examination of synteny within the *Nr2f* genomic environments. With respect to the more ancient *Nr2f1/2/5/6* genes, we did not find evidence of synteny between placozoans and cnidarians. However, the location of the three *N. vectensis* genes indicates they were likely derived from an ancient duplication event followed by a tandem duplication (S2 Fig). With respect to *Nr2f1/2/5* genes in invertebrates, we only found preservation of the *Nr2f1/2/5* loci between two slow-evolving deuterostomes: the amphioxus (*B. belcheri)* [43] and the hemichordate (*S. kowalevskii)* [44] (S2 Fig), corroborating the existence of the Nr2f1/2/5 cluster shown in the phylogenetic trees (Fig 2; S1 Fig). The only remaining synteny between *Nr2f1/2/5* invertebrates and vertebrate orthologs appears to be the linkage between *UNC45A*-*NR2F2* of primates and *Unc45a*-*Nr2f1/2/5* of the tunicate *C. robusta* (S3 Fig), which is considered the closest living relative of vertebrates [45]. Focusing on *Nr2f1* and *Nr2f2* in the genomes of gnathostomes, including cartilaginous fish, coelacanth, spotted gar, zebrafish, chicken, and human, we found a high degree of synteny for *Nr2f1* and *Nr2f2* loci and conservation of the location of flanking genes among these taxa (Fig 5). Specifically, *Nr2f1* and *Nr2f2* genes exhibited a remarkably conserved syntenic environment, clustering with putative orthologs belonging to other families. In fact, *Lysmd3, Arrdc3, Mctp1* and *Mef2a* flank *Nr2f1* orthologs, while *Nr2f2* orthologs are flanked by *Lysmd4*, *Arrdc4*, *Mctp2* and *Mef2c* paralogs. Furthermore, in teleosts like zebrafish, two *Nr2f1* Ohnologs (*nr2f1a* and *nr2f1b*) also shared significant conservation of paralogous genes, which is consistent with an origin from the teleost-specific genome duplication (TSGD) [15, 16]. However, the *nr2f1b* gene has been lost by several teleost species (Fig 2; S1 Fig). Although somewhat fragmented, orthologs of flanking genes found in gnathostome *Nr2f1* and *Nr2f2*, such as *Arrdc2/3*, *Lysmd3*, *Fam172a,* were also found near each of the three lamprey *Nr2f1/2* genes (S4 Fig), which is consistent with these genes arising from genome duplication(s) within the agnathan lineage [46]. Together, these results suggest that *Nr2f1* and *Nr2f2* of gnathostomes have a common origin and are derived from a WGD event [47, 48].

**Fig 5.**
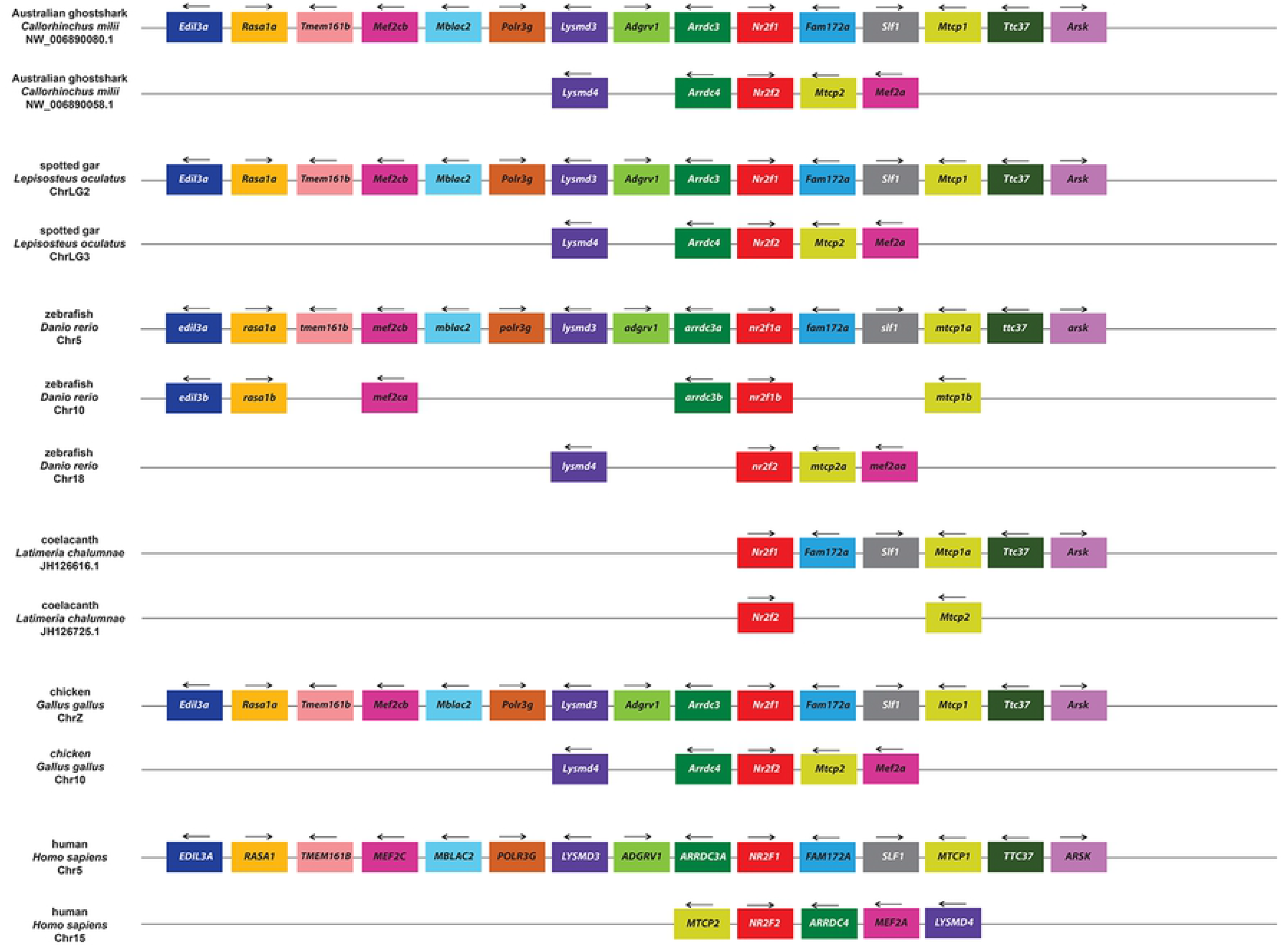
Synteny analysis of *Nr2f1* and *Nr2f2* genes. Schematization of conserved genome environments of gnathostome *Nr2f1* and *Nr2f2* genes (red rectangles) in selected species with relative chromosomes/scaffolds. Flanking orthologous genes are represented employing rectangles of the same color. Black arrows indicate transcription orientation.

Examining *Nr2f5* loci in the same representative gnathostomes showed a high degree of conservation in species that have retained the gene. However, the synteny is not shared with gnathostome *Nr2f1* and *Nr2f2* orthologs (Fig 6). The *Nr2f5* loci in coelacanth and amphibians have retained *Txnip* (Fig 6), which is also named *Arrdc6*. *Arrdc* family members flank both *Nr2f1* and *Nr2f2* genes (Fig 5). Interestingly, despite having lost *Nr2f5*, amniotes have largely preserved the flanking genomic loci present in fish in amphibians (Fig 6). In contrast, the absence of *Nr2f5* in cartilaginous fish, such as *C. milii*, correlates with the lack of the entire locus. Within the *Actinopterygii* (ray-finned fishes), the synteny of genes has been lost only on one side of the *Nr2f5* loci. With respect to the lamprey, its *Nr2f1/2C* ortholog is flanked by a *Bola1* ortholog as well as orthologs of gnathostome *Nr2f1* and *Nr2f2* (S4 Fig; Fig 5; Fig 6), which suggests ancestral linkage with the single *Nr2f1/2/5* genes (Fig 2; S1 Fig). As might be expected, the *Nr2f6* subfamily did not share common elements with the other *Nr2f* loci in gnathostomes (Fig 7). However, within gnathostomes the *Nr2f6* loci were highly conserved from cartilaginous fish to mammals, although there were significant gene losses surrounding *Nr2f6* loci in zebrafish and human. Furthermore, the presence of conserved orthologs (*ano8a* and *ano8b*, *plvapa* and *plvapb*) flanking *nr2f6a* and *nr2f6b* zebrafish genes suggested that they originated from the TSGD. Therefore, these findings show that despite the divergence of the *Nr2f5* and *Nr2f6* within vertebrates the genomic environments surrounding *Nr2f5* and *Nr2f6* loci are highly conserved within gnathostomes.

**Fig 6.**
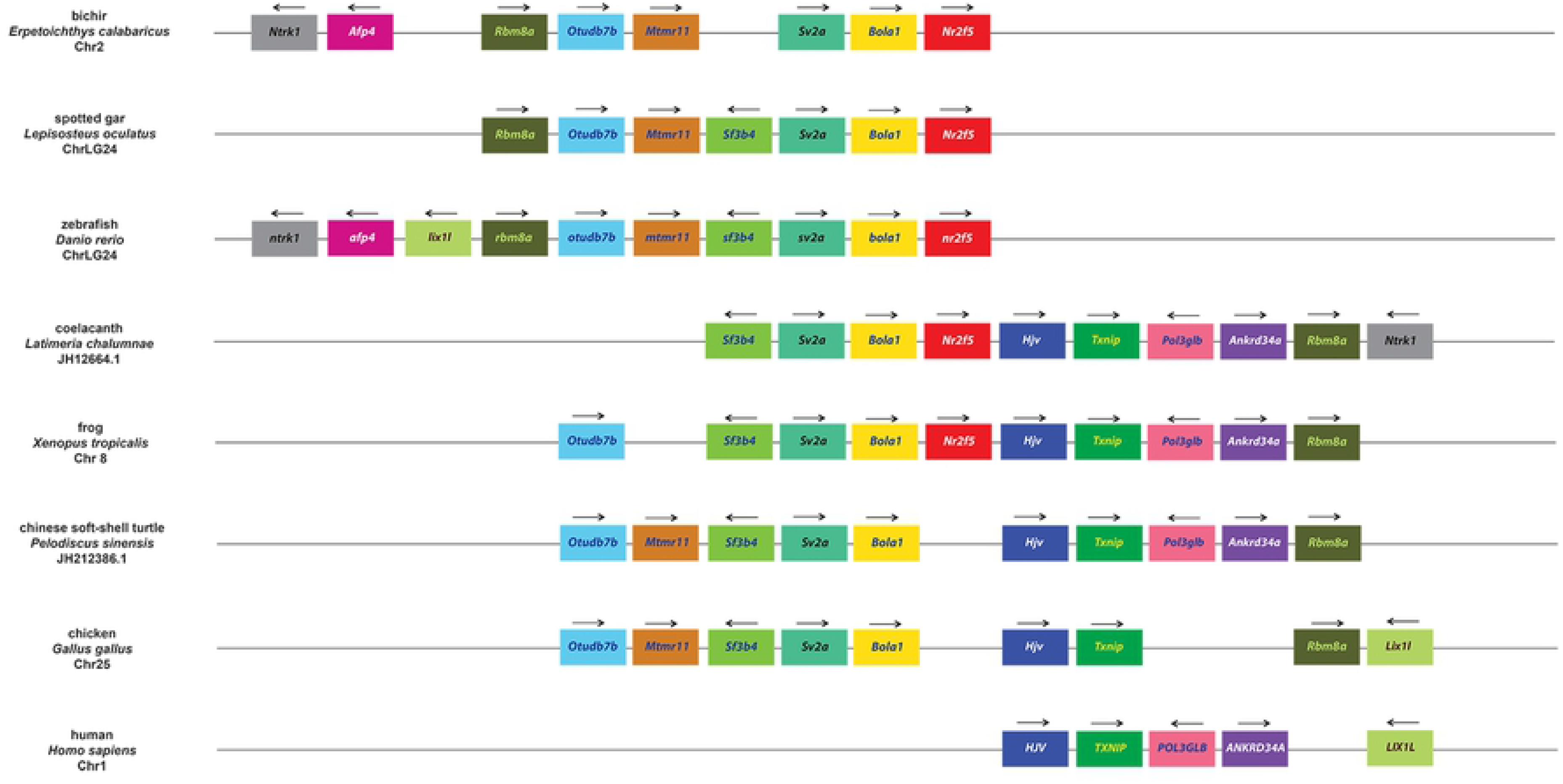
Synteny analysis of *Nr2f5* genes. Schematization of conserved genome environments of gnathostome *Nr2f5* genes (red rectangles) in selected species with relative chromosomes/scaffolds. Flanking orthologous genes are represented using rectangles of the same color. Black arrows indicate transcription orientation.

**Fig 7.**
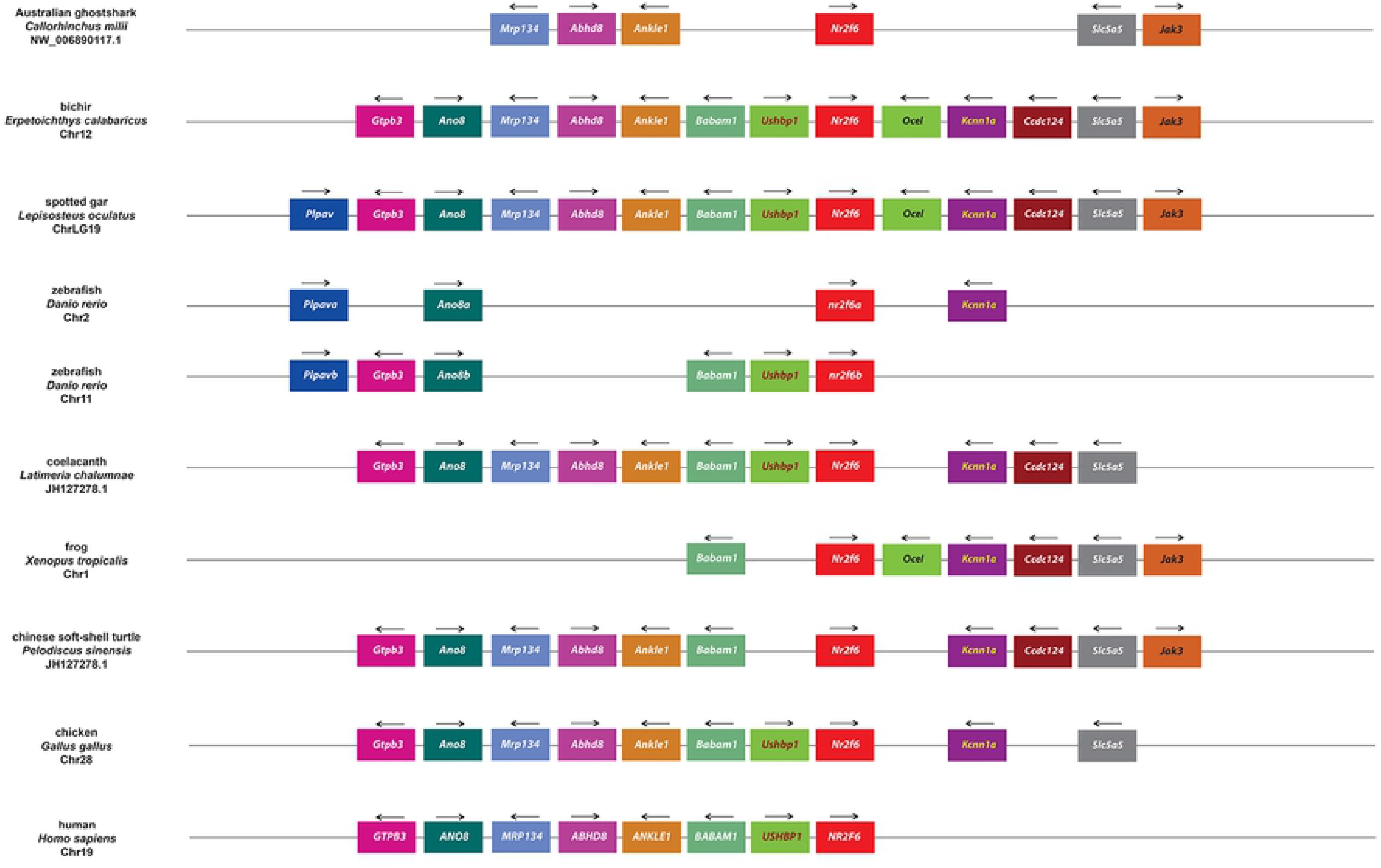
Synteny analysis of *Nr2f6* genes. Schematization of conserved genomic environments of gnathostome *Nr2f5* genes (red rectangles) in selected species with relative chromosomes/scaffolds. Flanking orthologous genes are represented using the same color code. Black arrows indicate transcription orientation.

### Effects of TSGD on the *Nr2f* gene repertoire

We next wanted to measure the impact of the series of additional WGDs that have occurred in teleosts on *Nr2f* gene number (Fig 2; S1 Fig). For this comparison, we examined all the *Nr2f* loci in zebrafish, the Asian arowana (*S. formousus*), which is documented to retain duplicates [49], and the Atlantic salmon (*S. salar*), which has a salmonid-specific genome duplication (SSGD) [39]. We found that each of these teleosts retained two *Nr2f1* Ohnologs (Fig 8), suggesting they either were not duplicated or that one pair of Ohnologs was lost in salmonids. Zebrafish lost one *nr2f2* Ohnolog, maintaining only the *nr2f2a* ortholog, while Asian arowana retained two *Nr2f2* Ohnologs. Salmonids have 3 *Nr2f2* genes (*Nr2f2a1*, *Nr2f2a2*, and *Nr2f2b1*), due to a loss of the one of the *Nr2f2b* Ohnologs following their additional WGD. With respect to *Nr2f5*, only Atlantic salmon showed two copies, implying these were generated during the SSGD event, as suggested by the presence of two *Nr2f5* Ohnologs in other salmonids (*Oncorhynchus spp*.). Finally, zebrafish and Asian arowana each possess two *Nr2f6* genes, while Atlantic salmon has 3 similar to what is found in the *Nr2f2* subfamily (Fig 8). Inspecting other teleost *Nr2f* gene family repertoires (Fig 2; S1 Fig), we found that the Channel catfish (*Ictalurus punctatus*), Red-bellied piranha (*Pygocentrus nattereri*), cavefish (*Astyanax mexicanus*) and Sheepshead minnow (*Cyprinodon variegatus*) all retained only *Nr2f6b*. The Sheepshead minnow and Princess cichlid (*Neolamprologus brichardi*) also lost *Nr2f5*. However, other cichlids like Blue tilapia (*Oreochromis aureus*) and Zebra mbuna (*Maylandia zebra*) did not lose *Nr2f5* (data not shown). Intriguingly, the Monterrey platifish (*Xiphophorus couchianus*) is the only gnathostome without any *Nr2f1* paralogs, differing from its sibling species, the common platifish (*X. maculatus*), which possesses *Nr2f1a*. Therefore, teleosts show an expansion of *Nr2f* genes following TSGD and SSGD, which were followed by high variability in species-specific losses of *Nr2f* Ohnologs.

**Fig 8.**
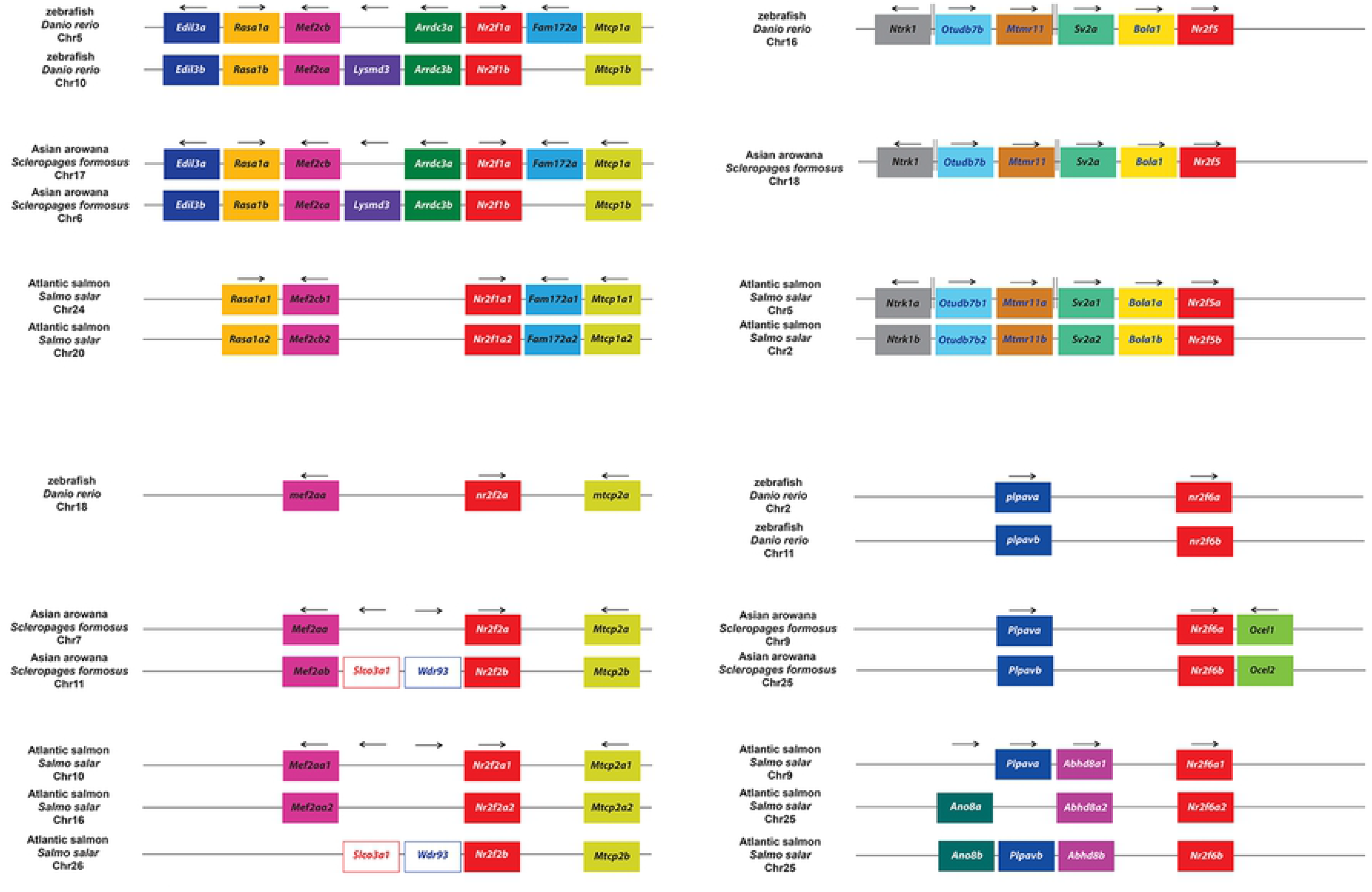
Synteny analysis of *Nr2f* genes in teleosts. Comparison of *Nr2f* genome environments in selected teleosts (zebrafish, Asian arowana, Atlantic salmon) with relative chromosomes/scaffolds. Rectangles of the same color represent flanking orthologous genes. Black arrows indicate transcription orientation.

## Discussion

We have performed an extensive examination of *Nr2f* gene evolution in metazoans. Our analysis corroborates previous work showing that *Nr2f* genes are present in some early-branching metazoans (cnidarians, placozoans) [13, 50], but that they are absent in other early-branching metazoans, i.e. sponges and ctenophores. Importantly, our data support a model in which a single *Nr2f* gene (*Nr2f1/2/5/6*), which is present in extant cnidarians and placozoans, predated the six Bilateria subfamilies that include *Nr2f1/2/5* (found in invertebrates), *Nr2f1/5* (found in agnathans), and *Nr2f1*, *Nr2f2*, *Nr2f5*, and *Nr2f6* (found in vertebrates; Fig 9). Single, conserved *Nr2f1/2/5* genes are predominantly found throughout invertebrate protostomes and deuterostomes and have even been retained in species traditionally considered gene losers, such as the tunicates [42,51,52]. There has been significant expansion and retention of *Nr2fs* in gnathostomes, particularly in teleosts. Although initial analysis in lampreys suggested they may possess only one *Nr2f* gene [53], our evolutionary assessment shows that the agnathans have three *Nr2f* members, which appear to have originated in part from an agnathan WGD event [46]. Interestingly, the single Nr2f1/2/5 proteins in invertebrates are also highly conserved at the sequence level and cluster with the Nr2f1/2 proteins in agnathans and Nr2f1 and Nr2f2 proteins in gnathostomes. Within gnathostomes, *Nr2f1* and *Nr2f2* orthologs have retained significant synteny of their loci, consistent with similar requirements shown across vertebrate models [14,18,19]. According to our data, *Nr2f1*, *Nr2f2*, and *Nr2f5* descended from the same ancestor, as limited synteny exists between them with members of the *Arrdc* family. However, *Nr2f6* orthologs do not share syntenic genomic loci with *Nr2f1*, *Nr2f2,* and *Nr2f5* genes. They have only retained synteny with their respective orthologs and have a distinct intron/exon structure. Together with the Nr2f6 proteins being the most divergent Nr2f with respect to other gnathostome Nr2fs, these data suggest the Nr2f6 origin predated the gnathostome radiation.

**Fig 9.**
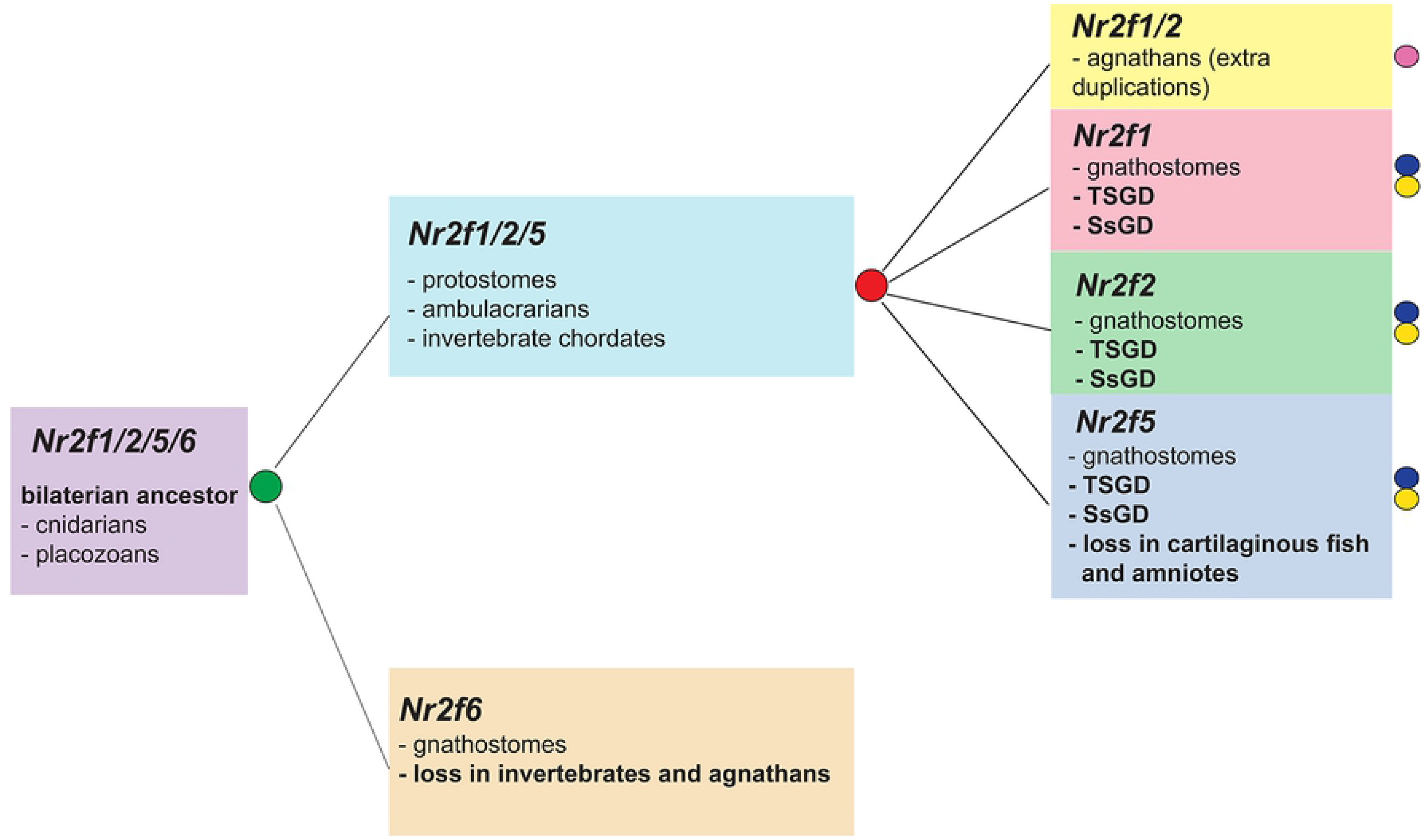
Evolutionary model for the *Nr2f* family in Metazoa. *Nr2f1/2/5/6* of placozoans and cnidarians (violet) is the ancestor of all other extant *Nr2fs*. An ancient duplicative event (green circle) generated *Nr2f1/2/5* (light blue) that is present from protostomes to invertebrate chordates and *Nr2f6* (orange), which has been retained only by gnathostomes and lost in invertebrates. WGDs in vertebrates (red circle) originated *Nr2f1/2* (yellow), *Nr2f1* (red), *Nr2f2* (green) and *Nr2f5* (blue), which has been lost in cartilaginous fish and amniotes. All these subfamilies show extra-duplicates: agnathan-specific (magenta circle) TSGDs (blue circle) and SSGDs (yellow circle).

Integrating our phylogenetic assessment with current expression and functional analyses of the *Nr2f1/2/5/*6 members in evolutionary distant animals [12, 54] supports a hypothesis that regulation of neuronal differentiation is the most ancient *Nr2f* function and that Nr2fs have gained additional functions in the development of mesodermal and endodermal-derived tissues. The *Nr2f* member (*Nr2f1/2/5/6*) of the diploblastic cnidarian *Hydra* only appears to be expressed in neurons and has a requirement in neurogenesis [12, 54]. Furthermore, the function of *Nr2f1/2/5* orthologs of protostome nematodes and flies have been extensively studied in neural sensory cell differentiation [8, 55], [56]. In invertebrate deuterostomes, the single *Nr2f1/2/5* orthologs are expressed in neural tissue of sea urchin (*Strongylocentrotus purpuratus*), amphioxus, and sea squirt embryos [11,57,58], [59]. Interestingly, recent functional analysis of the Mediterranean sea urchin (*Paracentrotus lividus*) Nr2f1/2/5 shows that it is maternally inherited and required for the development of neural and ectodermal derivatives [60]. A *Nr2f1/2/5* ortholog from the agnathan river lamprey (*Lampetra fluviatilis*) is also expressed in the developing nervous system [53]. However, our identification of three *Nr2f1/2* members in agnathans suggests that additional expression and potentially functional analysis should be performed in the sea lamprey (*Petromyzon marinus*) and/or hagfish (*Eptatretus burgeri*) to understand the conservation of the different agnathan paralogs compared to Nr2fs in vertebrates. In addition to the conservation of *Nr2f1/2/5/6, Nr2f1/2/5,* and *Nr2f1/2* expression in neural tissues from cnidarians to agnathans, *Nr2f* genes have also evolved critical new expression and functionalities, such as copulation control in nematodes [61] or heart vessels specification in flies [20, 21]. The recent work with the Mediterranean sea urchin suggests that it is required for the development of mesendodermal derivative [60], supporting that ancestral *Nr2fs* also acquired requirements in all 3 germ layers within the invertebrate lineage. However, the conservation of these requirements in the different germ layers throughout different metazoan phyla is not yet as clear, with present analyses suggesting functions in mesendodermal tissues are newly derived functions.

Similar to the expression in invertebrates, all the vertebrate *Nr2f* genes are expressed in neural tissue during development. Furthermore, it is interesting to consider the conservation of expression and functions of *Nr2f1* and *Nr2f2* genes in vertebrates compared to the analysis of the invertebrate *Nr2f* genes, since the functions of *Nr2f1* and *Nr2f2* have been intensely investigated in vertebrates and they are required for proper human development [19,28,62]. Our evolutionary analysis strongly supports paralogy between *Nr2f1* and *Nr2f2* of gnathostomes that is derived from one of the two rounds of WGD events in the vertebrate lineage [47, 48]. Consistent with the common origin for *Nr2f1* and *Nr2f2* genes in vertebrates, they share overlapping expression patterns and functions, but also appear to have acquired distinct developmental roles during evolution. Both *Nr2f1* and *Nr2f2* orthologs share overlapping central nervous system (CNS) expression in mouse and zebrafish [14,18,63], further supporting the conserved expression of *Nr2f* genes throughout neural tissue of metazoans. However, *nr2f1a* and *nr2f2* are both expressed more extensively in neural tissue of zebrafish embryos, following overlapping expression in early mouse embryos *Nr2f1* and *Nr2f2* become predominantly expressed in neural and mesendodermal tissues, respectively. Functional analysis of these *Nr2f* genes in mice and zebrafish support the functional divergence of these proteins. Murine *Nr2f1* KOs have glial differentiation defects [64], while *Nr2f2* is required for proper development of many mesendodermal-derived tissues, including atrial chamber and arterial-venous differentiation [37, 65]. Intriguingly, mouse *Nr2f2* and zebrafish *nr2f1a* are functional homologs with respect to heart development, as both are required for atrial differentiation [66], further supporting the common evolutionary origins of these paralogs. While zebrafish *nr2f2* is not required for early atrial or vein development [26], NR2F1 and NR2F2 proteins do appear to have redundant requirements, for instance promoting atrial cardiomyocytes differentiation in human embryonic stem cells [67, 68]. It is interesting that the single *Nr2f1/2/5* homolog of flies (*svp*) is also required for dorsal vessel (heart) development [20]. However, if these similar roles in heart tissues reflect conserved requirements within metazoans for cardiac differentiation requires functional studies from many additional model organisms, though the analysis in sea urchins hints at conservation of more ancestral requirements in all germ layers [60]. With respect to the expansion of *Nr2f1* and *Nr2f2* in teleosts*, Nr2f1b* has been lost in the majority of surveyed teleosts. *Nr2f1b* zebrafish mutants are viable [69] and do not exhibit redundancy with *nr2f1a* in atrial cardiomyocyte differentiation [26], but do exhibit some redundancy with multiple other *Nr2f* genes in neural crest cells that promote jaw development [69]. Virtually all the analyzed gnathostome genomes have a single *Nr2f2* gene, excluding the teleosts *S. formosus* (2) and *S. salar* (3), implying there may be some dosage sensitivity that favors the retention of single orthologs in vertebrates. Otherwise, the presence of additional duplicates in these species could suggest new functionalities.

With respect to *Nr2f5* orthologs, our analysis shows they are the smallest *Nr2f* subfamily in vertebrates. It has been maintained in coelacanth, ray-finned fish, and amphibians, but independently lost in cartilaginous fishes and mammals (Fig 9). Despite the loss of *Nr2f5*, the synteny of organisms that both have and have not retained *Nr2f5* suggests a common evolutionary origin with *Nr2f1* and *Nr2f2* paralogs. Similar to invertebrate *Nr2f* genes, *Nr2f1* and *Nr2f2* orthologs, *Nr2f5* is expressed in neural tissue [63], including in the eyes of zebrafish and newts [70, 71]. Like *nr2f1b*, zebrafish *nr2f5* mutants are viable, yet function redundantly with other *nr2f* genes for a proper upper-jaw development [69]. While expression and functional analysis from other organisms that have retained *Nr2f5* (coelacanth, spotted gar, and frog) may provide insights into conserved functions of *Nr2f5* orthologs, the loss of *Nr2f5* lineage in cartilaginous fish and amniotes, as well as the lack of overt requirements alone in zebrafish, suggests that *Nr2f5* orthologs likely have retained singular developmental requirements within the vertebrate lineage.

In contrast to *Nr2f5*, the divergent *Nr2f6* subfamily has been retained by all the evaluated gnathostomes and exhibit high sequence conservation, suggesting pressure to maintain these genes in vertebrates. Intriguingly, the high degree of divergence of the *Nr2f6* subfamily evokes a pre-gnathostome origin for this gene, as has been proposed from more limited analysis [72]. Despite the divergence to other *Nr2f* subfamilies, *Nr2f6* genes have conserved expression within the central nervous system of mammals [14], as well as both zebrafish *nr2f6* Ohnologs. Murine *Nr2f6* KO mice have forebrain defects. Specifically, a loss of neurons that regulate the circadian clock genes [73]. *Nr2f6* also has a critical role in lymphocyte differentiation and T-cell mediate tumor surveillance, suggesting derived functions in adaptive immunity [74, 75]. Thus, our data support a pre-gnathostome origin for the *Nr2f6* subfamily during metazoan evolution and that it has been selectively retained in gnathostomes.

In examining the evolution of the Nr2f transcription factors, it is also worthwhile to note that there is conserved responsiveness to retinoic acid (RA) signaling that exists in early-branching metazoans through invertebrate chordates and gnathostomes, implying this relationship may form the core of an ancient gene regulatory network. *Nr2f* genes from placozoans [50] and the invertebrate chordates *Ciona* and amphioxus are all RA-responsive [11, 57]. Furthermore, in vertebrates, where the earliest requirement for RA is posteriorization of the embryo [76], virtually all the *Nr2f* genes have been shown to be responsive to RA signaling in developmental contexts involving all three germ layers. Specifically, RA signaling has been shown to positively regulate all the *Nr2fs* in zebrafish in developing zebrafish endoderm [77], the CNS [63], and anterior lateral plate mesoderm (ALPM) [26]. RA signaling also positively regulates *Nr2f1*, *Nr2f2* and *Nr2f6* in mice [78, 79], and *NR2F1* and *NR2F2* in humans [80, 81]. Nr2fs can inhibit RA signaling in some contexts, suggesting it may form a negative feedback loop. One role Nr2fs may play is through direct competition with retinoic acid receptors (RARs) in binding retinoic acid response elements (RAREs) [18]. Moreover, it has been shown that the cnidarian Nr2f1/2/5/6 possesses the ability to inhibit RA signaling in *in vitro* signaling assays [12]. Thus, the responsiveness of the *Nr2f* family to RA may have evolved very early and has been highly maintained through the diversification of multiple vertebrate *Nr2f* genes, implying there is high selection to maintain this relationship.

## Conclusions

Overall, our evolutionary assessment sheds new light on the events that have shaped the extant *Nr2f* family in Metazoa. The phylogenetic analysis clearly defines the individual *Nr2f* subfamilies and their relationships across metazoan phyla, which complements available expression and functional data that together support an origin of their requirements in the development of neural tissue. Interestingly, the functions of Nr2f proteins are found to regulate development of virtually all germ layers of invertebrates and vertebrates. The detailed evolutionary understanding of the *Nr2f* gene family we now have will allow us to infer more meaningful conclusions about the origins and conserved requirements of *Nr2f* genes in normal metazoan development and their role in sculpting diverse body plans.

## Methods

### Ethics Statement

Ethical approval is not required. No animals were used in this study.

### Genome database searches and phylogenetic reconstruction

*Homo sapiens* NR2F protein sequences were employed as queries in BLASTp and tBLASTn in genome databases of selected species (NCBI, Ensembl, Ensembl Metazoa, SkateBase [82], ANISEED [83]). The entire dataset of protein sequences for domain architecture was analyzed by using the domain database provided by Expasy, named PROSITE [84] and then, manually annotated. All the surveyed sequences were verified to be Nr2f proteins through analysis of DBDs and LBDs (S2 File). The analysis was weighted with 30 species from agnathans to primates to take into account the impact of multiple WGDs in vertebrates [47, 48] and in teleosts [15, 16]. Orthology of the *Nr2f* members was initially assessed by using a reciprocal best blast hit (RBBH) approach employing default parameters and corroborated by phylogenetic analyses. Protein alignment for phylogeny was generated using L-INS-i (accurate; for alignment of <200 sequences) on MAFFT [85, 86] (S3 File). The phylogenetic reconstruction of Fig 2 was performed on the entire protein sequences and based on maximum-likelihood (ML) inferences calculated with PhyML 3.0 [87], employing automatic Akaike Information Criterion (AIC) by Smart Model Substitution (SMS) [88], which selected the JTT+G+F model employing discrete gamma distribution in categories. All parameters (gamma shape = 0.692; proportion of invariants (fixed) = 0.000) were established from the dataset. Branch support was provided by aLRT [89]. The phylogeny of S1 Fig was carried out employing Bayesian Information Criterion (BIC) by SMS, which sorted the JTT+G+F model using discrete gamma distribution in categories. All parameters (gamma shape = 0.690; proportion of invariants (fixed) = 0.000) were established from the dataset, with branch support calculated employing aBayes method [90]. Accession numbers and protein sequences used for phylogenetic tree reconstructions are provided in S4 File, while those excluded for their divergence are listed in S5 File. Common and Latin names for species used in this study are listed in S6 File.

### Analysis of intron/exon structures and phases

Gene structures were deduced after merging the genomic sequences with ESTs when available, as previously described [40–42]. Introns were classified as phase 0, phase 1, and phase 2, according to their positions with respect to the protein-reading frame. The amino-acid residues with the conserved introns were manually mapped on a ClustalX alignment [91] of selected Nr2f proteins (S7 File).

### Evaluation of synteny

We evaluated the presence/absence of synteny examining the chromosomes on public genome databases (NCBI, Ensembl, Ensembl Metazoa, ANISEED [83]). We verified the existence of duplicates using Genomicus [92] and Vertebrate Ohnologs [93]. The window considered for the locus analyses was twenty flanking genes. Genes that were not conserved were excluded from the analysis. All the genes were represented employing colored rectangles, using the same color for all *Nr2f* genes (red).

## Acknowledgements

Not applicable

## Availability of Data and Materials

All the data used for the analysis in this article are available within the article and in its online supplementary material.

## Authors’ contribution

UC performed the analyses, prepared the figures and wrote the initial manuscript. JSW provided feedback for the analyses, revised figures, and wrote the manuscript.

## Supporting Information

**S1 Fig. Phylogenetic tree of the Nr2f family, using Bayesian Information Criterion (BIC).** The same color code as Fig 2 is used. Values at the branches indicate replicates obtained employing the aBayes method.

**S2 Fig. Synteny analysis of *Nr2f* genes in invertebrates.**

**A**) Schematic of *Nr2f1/2/5/6* gene duplications in *N. vectensis*. **B**) Schematic of conservation of *Nr2f1/2/5* loci between *S. kowalevskii* and *B. belcheri*. Black arrows indicate transcription orientation.

**S3 Fig. *Nr2f-Unc45* gene duplet preservation.**

Schematic of *Nr2f-Unc45* duplet conservation in genomes of ascidians (*Ciona*) and primates. The duplet is absent in other vertebrate models, including zebrafish and mouse.

**S4 Fig. Synteny analysis of *Nr2f1/2* lamprey genes.**

Schematic of lamprey (*P. marinus) Nr2f1/2* loci with relative chromosomes. Same color code of Figs. 3-6 is used. Flanking genes are in common with gnathostomes, with *Arrdc2* and -*3* (green) that form a conserved duplet with *Nr2f1/2B* and *Nr2f1/2C*. Black arrows indicate transcription orientation.

**S1 File. (docx) Intron/exon structure of *Nr2f* genes in Metazoa.**

Alignment of specific and conserved intron/exon boundaries within the Zinc finger motifs of DBD (underlined) and LBDs (red). The intron phases have been depicted using color code: Phase 0 (yellow), Phase 1 (green), Phase 2 (turquoise).

**S2 File. (docx) Nr2f domain architectures during metazoan evolution.**

Sequences used in analysis with DBDs (yellow) and LBDs (green) domains in metazoan Nr2f proteins indicated. the Zinc-finger motifs within the DBDs are underlined.

**S3 File. (txt) MAFFT alignment of protein sequences used for phylogenetic analysis of** Fig 2 **and S1 Fig.**

**S4 File. (txt) List of all protein sequences employed in Nr2f phylogenetic tree with accession numbers.**

**S5 File. (txt) List of protein sequences excluded from Nr2f phylogenetic tree due to their high degree of divergence.**

**S6 File. (xlsx) List of species used for our evolutionary analyses with their common names, Latin names, and phyla.**

**S7 File. (docx) Selected Nr2f transcripts and translations used for analysis with the positions and phases of intron/exon boundaries indicated.**

